# The role of host cell glycans on virus infectivity: The SARS-CoV-2 case

**DOI:** 10.1101/2021.05.08.443212

**Authors:** Silvia Acosta-Gutiérrez, Joseph Buckley, Giuseppe Battaglia

## Abstract

Long and complex chains of sugars, called glycans, often coat both the cell and protein surface. Glycans both modulate specific interactions and protect cells. On the cell surface, these sugars form a cushion known as the glycocalyx. Here, we show that Heparan Sulfate (HS) chains – part of the glycocalyx – and other glycans – expressed on the surface of both host and virus proteins – have a critical role in modulating both attractive and repulsive potentials during viral infection. We analyse the SARS-CoV-2 virus, modelling its spike proteins binding to HS chains and two key entry receptors, ACE2 and TMPRSS2. We include the volume exclusion effect imposed on the HS chains impose during virus insertion into glycocalyx and the steric repulsion caused by changes in the conformation of the ACE2 glycans involved in binding to the spike. We then combine all these interactions, showing that the interplay of all these components is critical to the behaviour of the virus. We show that the virus tropism depends on the combinatorial expression of both HS chains and receptors. Finally, we demonstrate that when both HS chains and entry receptors express at high density, steric effects dominate the interaction, preventing infection.

## Introduction

The COVID-19 pandemic, caused by the Severe Acute Respiratory Syndrome Coronavirus 2 (SARS-CoV-2), has caused over 124 million infections and over 3 million deaths worldwide (World Health Organisation, WHO). SARS-CoV-2 belongs to the betacoronavirus genus of the coronaviridae family, the same as the Middle East Respiratory Syndrome coronavirus (MERS-CoV), and the Severe Acute Respiratory Syndrome coronavirus (SARS-CoV), two well established pathogens which emerged in the middle east in 2012 and the south of China in 2003, respectively [1] These outbreaks have demonstrated that coronaviruses have a high zoonotic potential, and therefore, understanding their mechanism of infection is necessary to improve our therapeutical approach and lessen the burden on our healthcare systems[2].

The virus life cycle involves a game of multivalency to enable binding to and infection of the host cell (**Figure 1**) [3]. Viruses recognise and bind to multiple cell surface receptors, either via protein/protein or protein/glycan interaction. Different viruses target different receptors – receptors which may have diverse sequences, structures, and cellular functions[4]. Viruses prefer the membrane proteins involved in cell adhesion [5]. Nature tightly regulates the single viral protein/receptor interaction or affinity and, moreover, their combination into multivalent and multiplexed association profiles which lead to sharp response to gradients of receptors density and give rise to the so-called super-selective binding [6].

**Figure 1:**
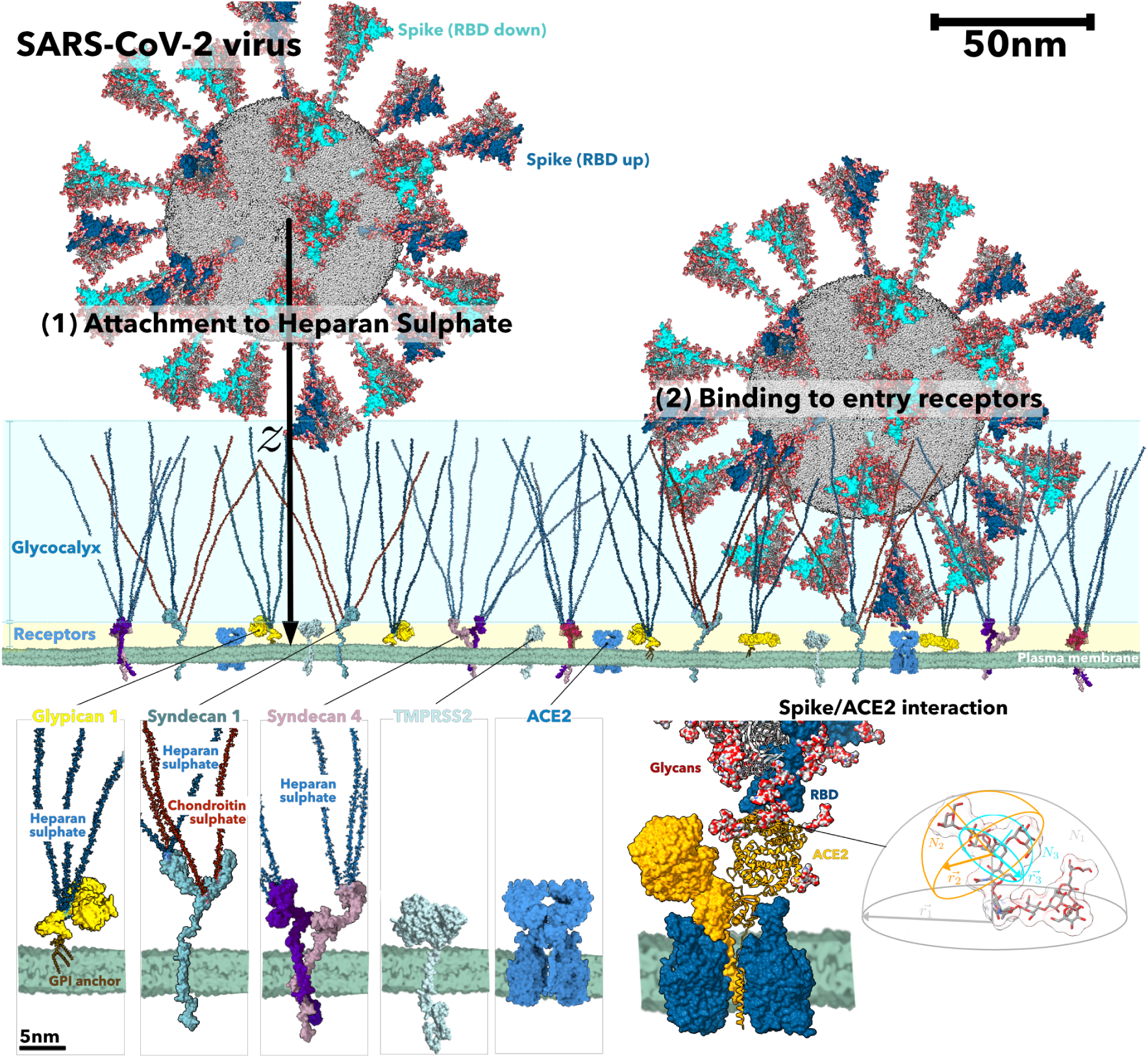
SARS-Cov 2 entry into the host cell. (a) Representation of the SARS-CoV-2 virion landing on the membrane of a host cell. SARS-CoV-2 spike proteins are shown as a surface with all-atom glycosylation. Protomers are coloured according to the RDB conformation: up (purple, PDBid 6VXX) or down (cyan, PDBid 6VSB). On the host membrane, all-atom structures of different components are represented as surface: Syndecan 1 and 4, Glypican, TMPRSS2 and the ACE2-BT0 complex (PDBid 6M1D). (b) Insertion into the glycocalyx: the insertion of one spike protein (in the closed state) into the glycocalyx. (c) Host receptor (ACE2) recognition and binding: the ACE2-BT0 complex structure bound to one spike protein in the open conformation. The binding region between one ACE2 monomer and the RBD domain of one protomer of the spike protein is highlighted as an inset. (d) Glycan conformational volume: an all-atom representation of a high-mannose glycan and its conformational volume.

Most human cells, particularly those that comprise biological barriers such as endothelial and epithelial cells, have a membrane coated by a dense and complex mixture of long polysaccharides, knows as glycosaminoglycan (GAG) chains. These chains bind to specific proteins forming compounds called proteoglycans. This chemical cushion, known as the glycocalyx, has several functions, including protecting cells from chemical injury and hindering the entry of pathogens. Several viruses have evolved to exploit these glycans, using them as an attachment factor to facilitate infection [7, 8]. SARS-CoV-2 is an example of this, binding to Heparan Sulfate (HS), one of the most abundant components of the glycocalyx [9, 10, 11]. Additionally, it has been shown that Heparin – a Heparan Sulfate analogue and widely used anticoagulant, antithrombotic, and blood-thinning drug – inhibits infection in-vitro [9, 12, 11]. Several clinical reports have also suggested the therapeutic effect of heparin in both moderate and severe COVID-19, and there are currently over 90 current clinical trials ongoing investigating this [13].

As shown in **Figure 1**, the SARS-CoV-2 virus spike glycoprotein interacts with the Heparan Sulfate (HS) chains expressed by proteoglycans[14]. There are two families of proteoglycans, Syndecans (from 1 to 4) and Glypicans (from 1 to 6); in **Figure 1**, we show the structure of the most common Syndecan-1 and −4 and Glypican-1. The affinity between the virus spike and the HS chains is dependent on the HS sequence[11]. Once the virus reaches the cell membrane, the specific binding of the receptor-binding-domain (RBD) of the spike glycoprotein (S) to the cellular-entry receptors (R) takes place (**Figure 1**). Different entry receptors have been identified for SARS-CoV-2, including angiotensin-converting enzyme 2 or ACE2 (HCoV-NL63, SARS-CoV and SARS-CoV-2) [15], TMPSRSS2, [15], SRB1 [16], neuropilin-1 [17]. The expression and tissue distribution of entry receptors influence viral tropism and pathogenicity, but the cell glycocalyx composition also plays a key role. In this paper, we present an analytical and predictive model for viral host tropism, based on the components of the cell-virus interaction during viral attachment.

### A simple model for Viral Tropism

The specificity of infection for certain viruses to hosts and host tissues, viral-host tropism, can be described in terms of the number of viruses bound to a given combination of receptors, or phenotype. The binding to the cell surface can be estimated using an Langmuir isotherm. We define two surface coverages, the glycocalyx surface coverage, *θ*_*G*_, which represents the fraction of the glycocalyx occupied by virions, and the receptor surface coverage, *θ*_*R*_, which represents the fraction of the cell-surface occupied by viruses,[18, 6, 19] equal to

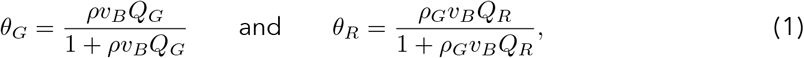

where *ρ* is the bulk viral titre (the number of virus copies per unit volume or viral load), *ρ*_*G*_ is the viral titre in the glycocalyx, *v*_*B*_ is the binding volume, and *Q*_*R*_ and *Q*_*G*_ are the grand canonical partition functions for the virus in the receptors and the glycocalyx surface, respectively. The surface coverage, *θ*, effectively normalises the virus binding: *θ* = 0 indicates no interaction and *θ* = 1 indicates full saturation of the available binding sites on the surface considered. Such an approach allows us to assess the nonlinear nature of viral binding and how it changes from zero to saturation for very small changes of the cell’s receptor/HS composition and the spike affinities towards them, viral size, spike number and initial viral load. The grand canonical partition functions, *Q*_*G*_ and *Q*_*R*_, can be broken down into the product of the component partition functions arising from all the possible binding and steric potentials acting on the virus. We can divide this into HS binding 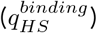, HS insertion 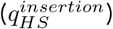, and receptor binding 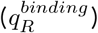 components respectively, as

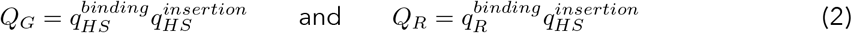

where

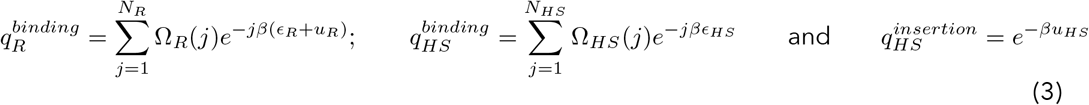

Here, *N*_*R*_ is the number of linkers, Ω_*R*_(*j*) the number of possible combinations between the R linkers and the virus proteins, *ϵ*_*R*_ the binding energy of the single bond between the virus protein and the R linker, *u*_*R*_ the R steric potential arising from compression of the receptor glycans (*u*_*R*_=0 for TMPRSS2). *N*_*HS*_ is number of HS chains per binding *µm*^2^, Ω_*HS*_(*j*) the number of possible combinations between the HS chains and the virus proteins, *ϵ*_*HS*_ the binding energy of the single bond between the virus protein and the HS chain, *u*_*HS*_, the steric potential arising from insertion of the virus into the glycocalyx, *β* is the inverse thermodynamic temperature. Equations (1-3) allow us to estimate the virus tropism as a function of the individual interactions, each of which can be derived separately and analysed down to the molecular level. Further derivation is provided in the Supporting Information.

As shown in **Figure 1**, during the insertion into the glycocalyx, coronaviruses bind to Heparan Sulfate. Each Heparan Sulfate chain, consists of a variably sulfated repeating disaccharide unit (**Figure 2a**). Only a fraction of these units bind to the virion spike proteins, normally 8 to 10 monomers or binding motifs [20]. Therefore, the total binding energy between a single HS-chain and the virus, *ϵ*_*HS*_, (Supporting Information) is dependent on the number of binding motifs, *N*_*L*_. The more binding motifs the higher the attractive energy (**Figure 2b**). This energy also increases with the number of HS chains, proportional to the density of proteoglycans (**Figure 2b**). The repulsive energy generated by the insertion of the virus into the HS, 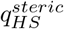, (Supporting Information) increases dramatically with the number of monomers per chain and the number of chains (proteoglycan density) (**Figure 2c**).

**Figure 2:**
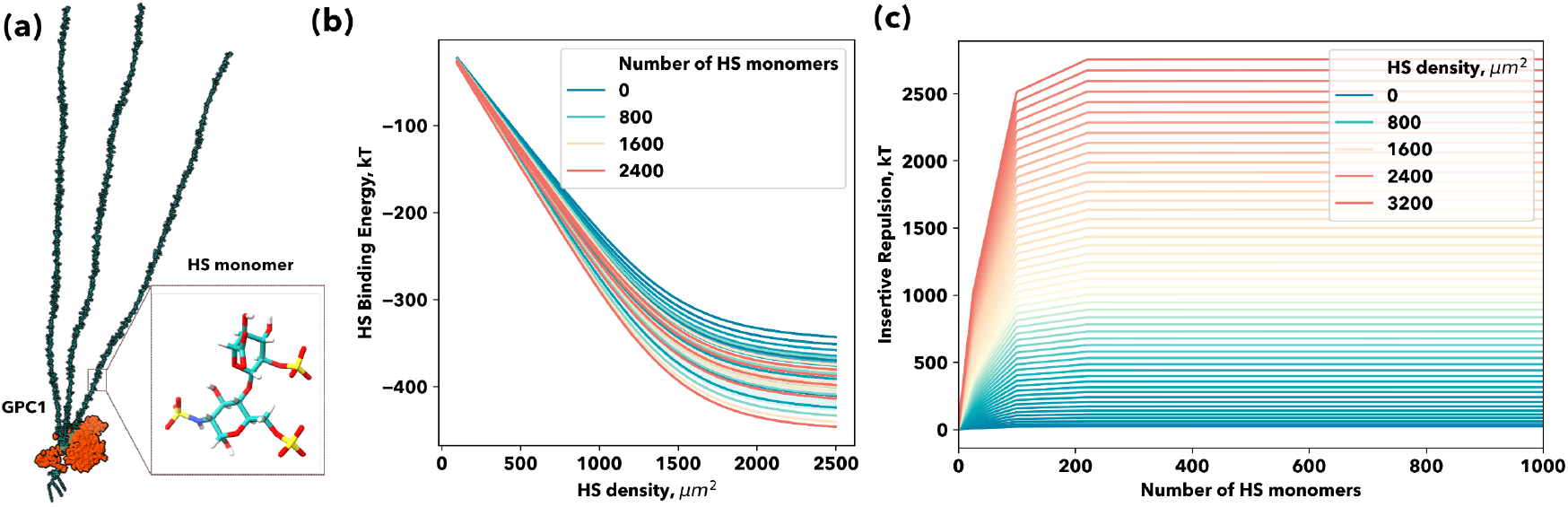
The insertion step into the glycocalyx. (a) All-atom representation of the glypican 1 proteoglycan together with an inset of an HS highly sulphated, IdoA(2S)-GlcNS(6S), monomer. (b) Heparan Sulfate (HS)-spike protein binding: binding interaction (kT) behaviour with increasing number of monomers per HS chain and HS chains density per *µm*^2^. (c) Heparan Sulfate (HS)-spike protein insertive repulsion: steric repulsion evolution with increasing number of monomers per HS chain and HS chains density per *µm*^2^.

Once a virus has inserted into the glycocalyx, it can bind to the ACE2 receptor via the RBD of the spike glycoprotein (**Figure 1**). Upon binding, glycans at the binding site are compressed, inducing a repulsive component to this interaction (**Figure 1** (lower right panel), **Figure 3a,b**, Supporting Information). The composition of these glycans can vary widely between species, with human glycans being particularly large and complex[21]. Both host-cell proteins and viral proteins are glycosylated, and this post-translational glycosylation process is known to play a major role in infection virulence and interaction with the host immune system [22]. In SARS-CoV-2, the spike protein typically contains 22 N-glycans [23, 24] and 2 O-glycans [23]. The spike protein glycans shield the virus from the immune system[25], but also seems to regulate the opening and closing of the RBD domain[26].

**Figure 3:**
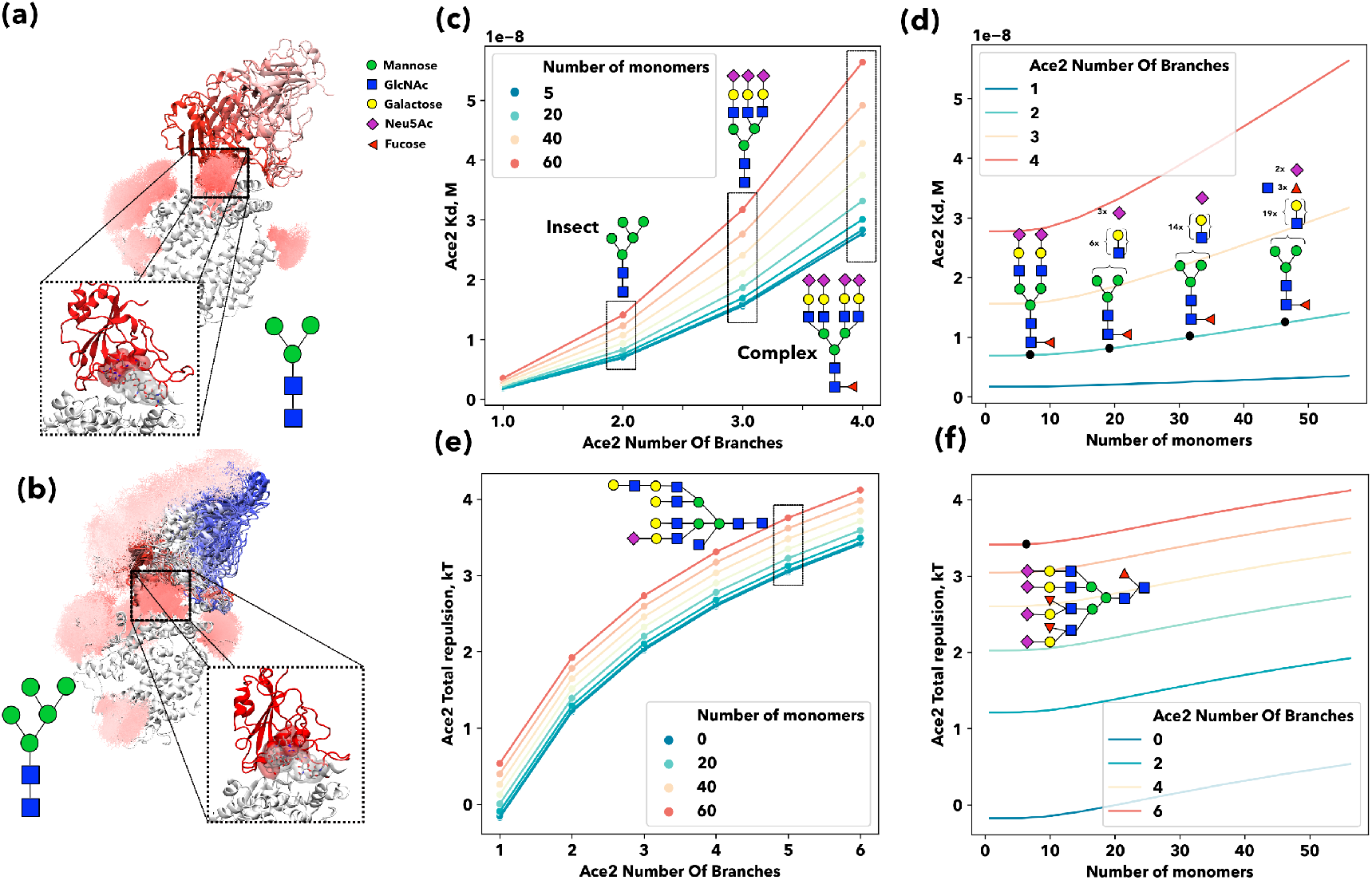
The effect of glycosylation in the recognition and binding of the spike protein to the host cell entry receptor.. (a, b) ACE2-RBD complex dissociation: multiple snapshots from an out-of-equilibrium simulation of the dissociation of the ACE2-RBD complex are super-imposed for the glycans and RBD domain. The RBD-domain is coloured according to the distance to the binding site (red bound, pink (a) or blue (b) unbound. Glycans are represented as lines and coloured radially in shades of pink. In both panels an schematic representation of the glycans is depicted as well as an all-atom representation of the glycan trapped in the binding site. (c,d) ACE2-RBD KD variation with glycan morphology: ACE2-RBD dissociation constant plotted as a function of the number of branches and coloured according to number of monomers per branch in c and vice versa in d. (e,f) ACE2 Glycans steric potential variation with glycan morphology: the steric potential generated by glycan compression in kT units is plotted as a function of the number of branches and coloured according to number of monomers per branch in e and vice versa in f.

In **Figure 3** we also analyse the influence of the trapped glycans on the dissociation constant (*K*_*D*_) of the ACE2-spike protein complex. Both the number of branches and branch length (number of monomers) decreases the *K*_*D*_ of the complex. Our analysis explains why the same spike proteins interacting with the same receptors are reported by several studies in human cells with different dissociation constants; proteins expressed in HEK293T-gave a *K*_*D*_ = 44.8 nM [27], FreeStyle293F *K*_*D*_ = 34.6 nM [28], while proteins expressed in insect Sf9 cells, reported *K*_*D*_ = 4nM [29]. Recently, Allen et al. [30] attempted to understand the influence of ACE2 glycosylation on the binding of the RBD domain of the SARS-CoV-2 spike protein. The authors chemically modified various glycan positions involved in the binding and measured the binding affinity using surface plasmon resonance (SPR). The predominant type of glycan observed across all five ACE2 glycosylated sites were fucosylated with two or three branches complex-type glycans which lack galactose and sialic acid. They found that ACE2 treated with sialidase has a higher affinity for SARS-CoV-2 compared to WT ACE2, KD 30% lower. The presence or absence of fucose or the presence of a single N-acetylglucosamine residue did not impact ACE2 binding to a significant extent. But when all N-linked glycans on ACE2 were converted to oligomannose-type glycans (predominantly Man9GlcNAc2) the *K*_*D*_ increases by 50%, corresponding to a decrease in binding. These differences, though subtle, are in perfect agreement with our model. In **Figure 3c,d** we observe that the increasing glycan complexity (branching and number of monomers per branch) increases the ACE2-RBD *K*_*D*_, due to the steric potential generated by the glycan compression in the binding site. The difference between an insect glycan (**Figure 3c**) and a more complex glycan (human-like) with 3 to 4 branches is nearly 20 nM (∼ 17.7 kT, for one glycan trapped in the binding region). According to our model, differences in *K*_*D*_ can be as high as 50 nM (∼ 16.8 kT) between a single-branched short glycan and multiple-branched long glycan, like the ones found in the human lungs (**Figure 3b**) [31]. Single chemical modification does not seem to have a major role in the complex binding affinity, as glycans have a steric role rather than a binding role. Therefore, the steric repulsion aids the dissociation of the complex – the shorter the glycan the higher the binding energy and the lower the *K*_*D*_. Addition-ally, we also performed out-of-equilibrium simulations of the RBD-ACE2 complex dissociation, using three different types of glycans: insect-like, high-mannose and complex. Our simulations reveal how the force needed to break the complexes decreases when the glycan complexity increases (**Figure S1**).

### Viral Tropism in a nutshell

The attractive/repulsive interactions described above defines the viral tropism, via the grand canonical partition function, *Q*_*HS*/*R*_. In our model, different cell phenotypes are characterised by the expression of proteoglycans (proteoglycan density, *µm*^2^) and the entry receptors (receptor density, *µm*^2^). Likewise, a virus can be described by the binding affinity of its spike protein to HS and the entry receptors. In **Figure 4a**, we see how the HS density and the virus spike-HS *K*_*D*_ affects *θ* for cells with a specific proteoglycan density, where above a certain proteoglycan density the virus-cell interaction is switched off – this is known as range selectivity[18]. This change from bound to unbound occurs within a very narrow range of HS densities. We can use the experimental data collected for SARS-CoV-2, SARS-CoV and MERS-CoV binding with Heparin [10], to show that SARS-CoV-2 has a higher affinity toward HS: this allows SARS-CoV-2 to bind to higher HS densities, compared to the other coronavirus (**Figure 4a**). By analysing the proteoglycan expression on the different tissues and cells found on the human upper and lower airways, **Figure 4b**, we found that the upper airways have a proteoglycan density which favours the attachment of SARS-CoV-2, but not SARS-CoV or MERS-CoV.

**Figure 4:**
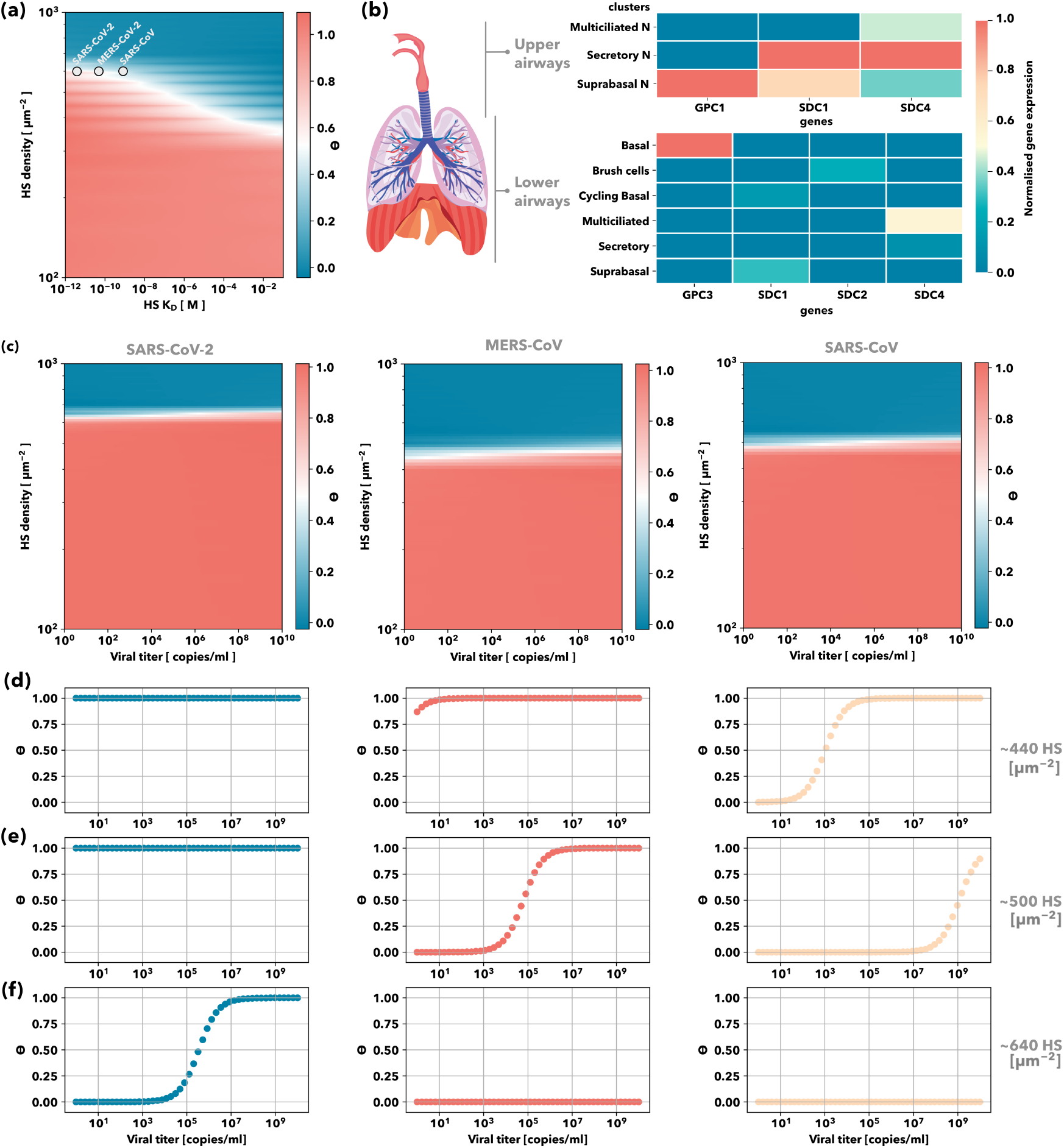
Viral tropism: cell proteoglycans density and initial viral titre effect. (a) Cell Surface coverage by viral particles variation with HS density and HS-spike binding affinity: Contour plot of the cell surface coverage by viral particle as a function of the HS density and the spike protein-HS binding affinity. Surface coverage equal to 1 implies that the initial viral titre successfully attaches to the cell surface. SARS-CoV-2, SARS-CoV and MERS-CoV spike proteins HS binding affinities are displayed. (b) Proteoglycans gene expression across cell types for the upper and lower human airways: Each gene is normalised to its maximum value across all tissues. The tissues correspond to the indicated positions along the airway from nasal to distal lung. Dataset obtained from Durant et al. Initial viral titre threshold for binding. (c) Cell surface coverage by viral particles variation with HS density and initial viral titre: Contour plot of the cell surface coverage by viral particle as a function of the HS density and the initial viral titre for SARS-CoV-2 (left), MERS-CoV (middle) and SARS-CoV (right). Cell surface coverage by viral particles variation with the initial viral titre for (d) ∼440, (e) ∼500 and (f) ∼640 HS chains per *µm*^2^; for SARS-CoV-2 (left), MERS-CoV (middle) and SARS-CoV (right).

The ideal location for attachment of MERS-CoV and SARS-CoV is in regions with lower proteoglycans densities, such as the lower airways, while SARS-CoV-2 can attach and replicate in the epithelium of the upper airways. This mean SARS-CoV-2 can infect the upper respiratory tract and can navigate the airways more easily, as reported by virological assessment of SARS-CoV-2, SARS-CoV and MERS-CoV patients [32, 33]. Moreover, we can observe in **Figure 4c** that the affinity of the virus for HS determines the initial viral titre require for successful attachment. At low HS densities, **Figure 4d**, very low initial titres result in full surface coverage for SARS-CoV-2 (**Figure 4d**, left) and MERS-CoV (**Figure 4d**, middle), while SARS-CoV needs at least 103 copies/ml in bulk to have 50% surface coverage. At medium HS densities, **Figure 4e**, SARS-CoV needs at least 109 copies/ml to achieve 50% surface coverage (**Figure 4e**, left) and MERS-CoV requires 105 copies/ml (**Figure 4e**, middle). At high HS densities, **Figure 4f**, MERS-CoV and SARS-CoV, cannot bind under any condition, while SARS-CoV-2 binds for initial viral titres above 10^3^ copies/ml. This is in agreement with respiratory samples extracted from SARS-CoV-2 patients with mild and severe COVID-19 symptoms revealed viral titres in between 10^3^ and 10^9^ copies/ml [12].

Several receptors have been found responsible for SARS-CoV-2 entry and while our analysis can be expanded to all of them, we focus on the most widely discussed elements in the SARS-CoV-2 infection, the receptors ACE2 and TMPRSS2[34]. In **Figure 5a** we observe how successful viral attachment occurs when the receptors act in a synergistic manner. Cells for which the number of ACE2 receptors account for less 40 % of the total receptor density per *µm*^2^, like the olfactory sensory neurons (OSN) [35, 36], have low cell surface coverage for any percentage of TMPRSS2, therefore, low viral titre on the surface. While for cell expressing high densities of ACE2, >60%, even moderate values of TMPRSS2 are enough to obtain high cell surface coverage values, as is the case for many cells from the human airways [37, 36], including olfactory epithelial sustentacular cells. This suggests that SARS-CoV-2 infection of non-neuronal cell types leads to olfactory dysfunction (anosmia) in patients with COVID-19.

**Figure 5:**
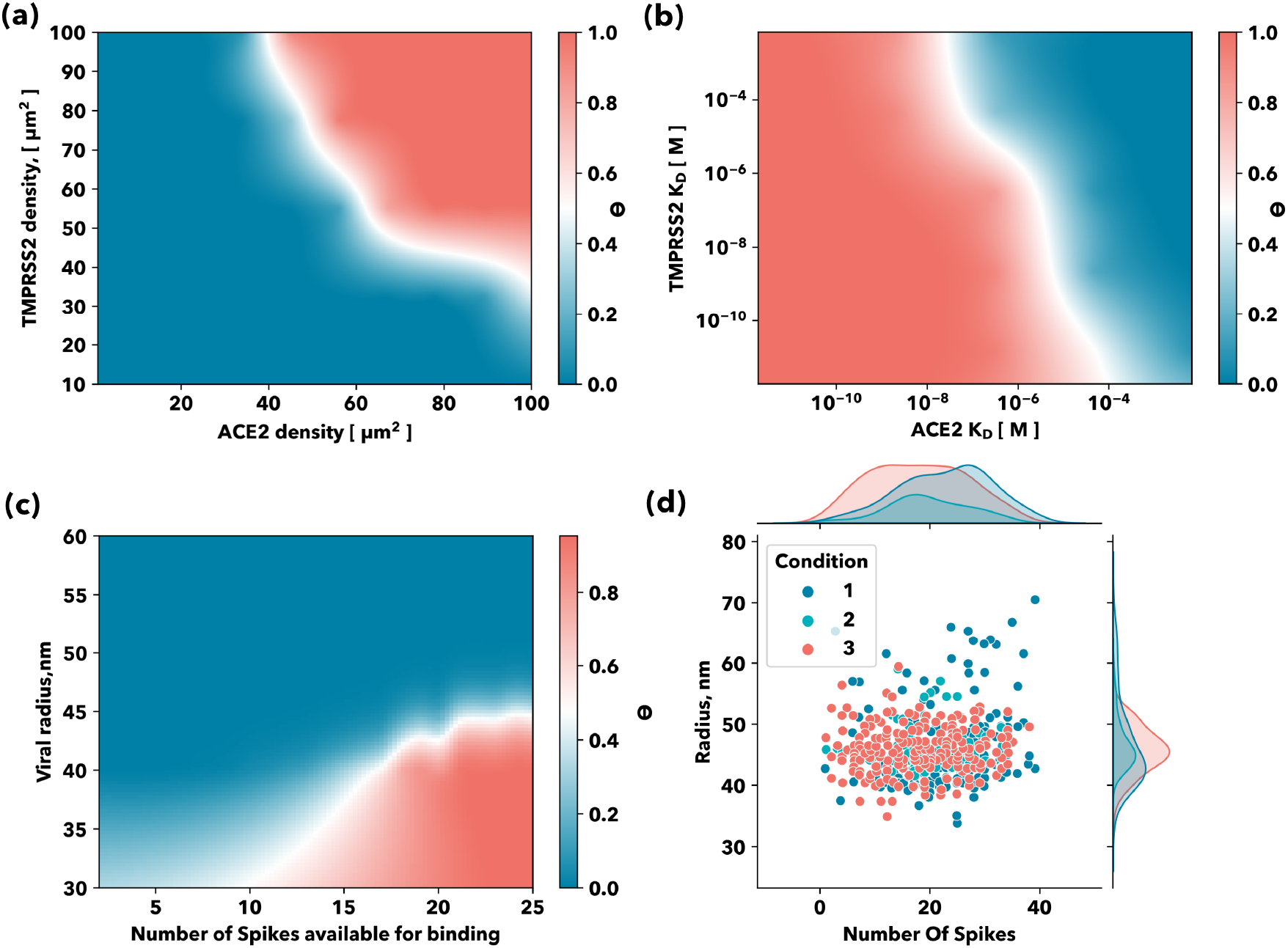
Viral Tropism: Entry receptors binding synergy. ((a) Contour plots of the cell surface coverage by viral particle as a function of the ACE2 and TMPRSS2 cell densities and (b) affinity towards the viral spike protein. (c) Viral morphology: radius and spike-proteins available for binding influence on viral the number of viral particles attached to the cell surface: Contour plot of the virus titre on the cell surface (Log) as a function of the virion radius and the number of spikes proteins in open conformation available for binding. (d) Viral morphology: radius and number of protein spikes. Probability distributions of the different SARS-CoV-2 viral radius and number of protein spikes. Data was replotted from Extended Data Fig. 1 in Ke, Z. et al. different sample conditions are reported on the original article [38].

Mutations of the RBD domain of the SARS-CoV-2 spike protein are worrying due to the higher transmissibility of these variants, especially the Brazilian (P1., K417T, E484K, N501Y), South African (B.1.351, K417N, E484K, N501Y) and British variants (B.1.1.7, N501Y) [39, 40, 41, 42, 43, 44]. This is related to the ability of these mutations to hide the virus from antibodies [39, 42, 44] and cause reinfection [41]. The N501Y mutation shared by all three variants also increases the binding affinity of the RBD-domain towards the ACE2, decreasing its *K*_*D*_[40]. However, for an optimum ratio of receptors (ACE2, TMPRSS2) densities, our model does not predict major changes in cell surface coverage with decreasing ACE2-RBD-domain *K*_*D*_ (**Figure 5b**). For high ACE2 *K*_*D*_ values (low binding affinity) on the other hand, changes in the affinity towards TMPRSS2 might lead to a decrease in viral titre on the cell surface according to our model (**Figure 5b**).

Finally, we found that both the virion morphology and the number of available RBD-domains in open conformation available for binding to the ACE2 affects virus titre on the cell surface. Smaller virions ∼40 nm radius are able to attach to the host cell with fewer spikes proteins available for binding to the entry receptors, while a bigger virus with same number of spikes available for binding would less successful in this case and the virus titre on the cell surface will be lower **Figure 5c**. Cryogenic electron microscopy on SARS-CoV-2 virions populations have revealed an average radius of ∼45 nm with a standard deviation of ∼10nm, with individual virions containing 24 ± 9 spike proteins not uniformly distributed and highly flexible [45, 38]. There is a very nonlinear response with the morphology of the virus, as shown in Figure 5c, and a virus with a radius larger than 45 nm is less effective at attaching to cell. According to our model, for larger virions to be as effective as viruses with a radius of 45 nm, they require more spike proteins on their surface. By fitting the measurements provided by Ke et al. [38](**Figure 5d**) we observe that for all three preparations there is a clear distribution of size, but the number of spikes per virion distributions are very broad, indicating that the size of the virus is the critical parameter for viral attachment.

## Conclusions

Overall, our model can explain the viral tropism of SARS-CoV-2 in terms of interactions with the various host-cell components. We find that viruses are extremely selective, not only in terms of entry receptors distribution on the host cell but also on the spike protein affinity towards the key elements of the cell phenotype. Additionally, we find that Heparan Sulfate plays a major role in viral infection, as the repulsion potential can dramatically decrease the number of viral particles that can interact with the entry receptors – this increases the viral load threshold required for infection. We have also shown that the role of the glycans on the entry receptors is not enthalpic but entropic, decreasing the binding affinity of the RBD-ACE2 complex at increasing morphological complexity. Finally, we have shown how the size of the viral particle is a critical parameter with a nonlinear response, and therefore, induces a selection of the optimal viral size for infection.

## Author contributions

J.B and G.B perform the mathematical derivation of the model. J.B. coded the model. S.A.G performed the all-atom simulations and the generation, analysis and representation of the synthetic data presented in the manuscript. All three authors wrote the manuscript.

## Competing interests

The authors declare no competing financial interests.

## Methods

### Out-of-equilibrium molecular dynamics

The ACE2-RDB SARS-CoV2 complex was extracted from the cryo-em pdb structure 6M17[46]. The protein complex presented no mutations or structural gaps. Two different systems (different glycan types) were generated by adding glycans using the Glycam web server Glycoprotein Builder tool[47], the insect-like system (DManpa1-3[DManpa1-6]DManpb1-4DGlcpNAcb1-4 DGlcpNAcb1-OH) and high mannose (DManpa1-6[DManpa1-3]DManpa1-6[DManpa1-3] DManpb1-4DGlcpNAcb1-4DGlcpNAcb1-OH). The two systems were protonated at pH 7.4 and solvated (150mM NaCl) using a TIP3P[48] water model and Charmm36m[49] protein, glycans and ions force field. The final simulated systems consisted of 240K atoms, resulting in a cubic box simulation cell of 12nmx12nmx17nm. All simulations were performed with Gromacs v2019.2 [50] with a 2fs integration step. We performed 2000 steps of minimization (steepest descent algorithm), followed by 5000 steps of isotropic NPT at 300K, using the velocity-rescaling thermostat[51] and Parrinello-Rahman barostat[52]. After 50000 steps of NVT the system was fully equilibrated and ready for production, which was performed in the NVT ensemble for 100 ns. A hundred steered molecular dynamics simulations were performed for each system using PLUMED2.6[53, 54].

### Synthetic data generation

In Table 1 we specify the model parameters fixed to generate the data for each of the figures displayed in this article.

**Table 1.**
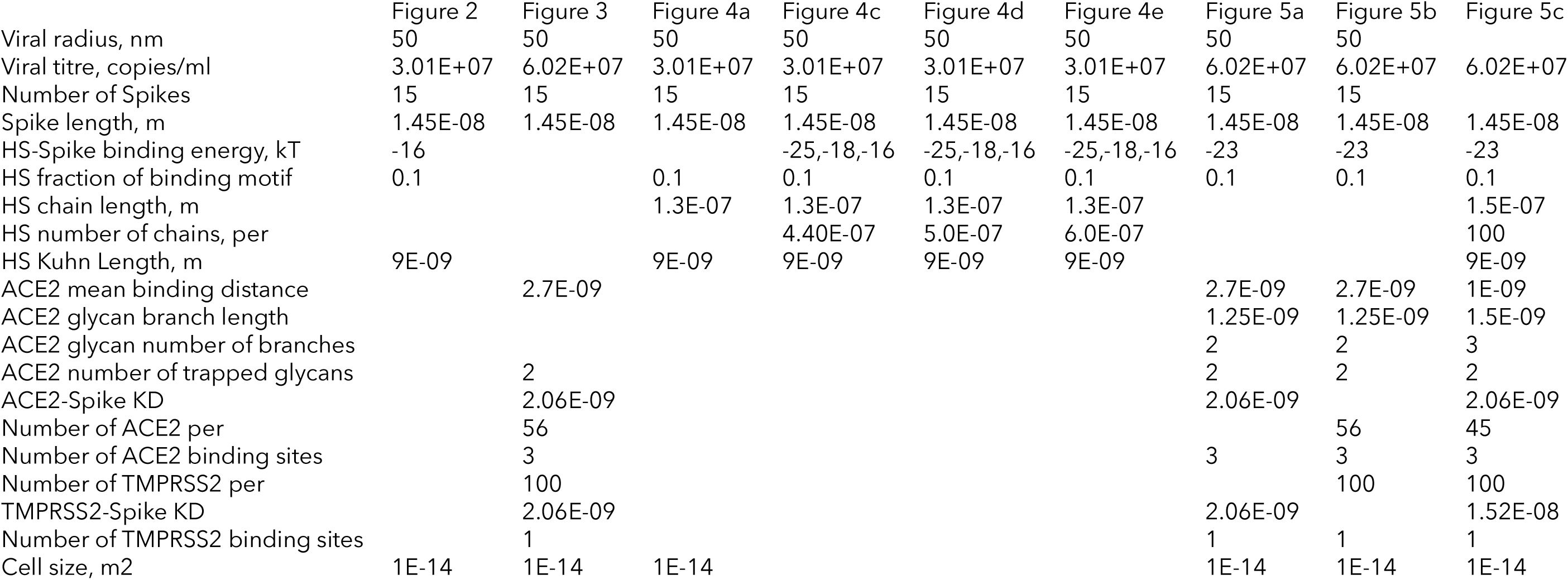
Model fixed parameters used to generate the synthetic data.

## Supporting information

### 1 Theoretical Model

As discussed in the main text, binding between coronaviruses and the host cells proceeds in a two step process. First, the virus binds to Heperan Sulfate, the long-chain sugars that coat the cell surface. After binding to the Heperan Sulfate, the virus then can form bonds with the receptors on the surface, particularly ACE2 and TMPRSS2.

We will define the glycocalyx surface coverage, *θ*_*G*_, as the probability that a virus will be bound to a HS chain at a given site, and the cell-surface coverage, *θ*_*R*_, as the probability that a virus will be bound to a receptor at a given site. In equation 1 in the main text, we give the general form of the surface coverage, derived from the Langmiur isotherm, which we will specialise to deal with each case, giving

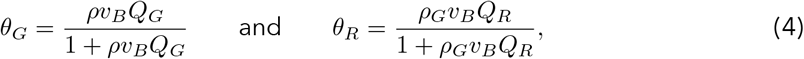

where the binding volume, *v*_*B*_, is determined from the viral radius, *R*, and the spike length, *d*, as 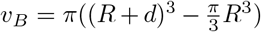 and *ρ* is the viral bulk concentration, and *ρ*_*G*_ is the viral concentration in the glycocalyx. The cell-surface coverage can be linked to the glycocalyx surface coverage as 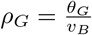, however, in order to limit the complexity for our calculation, we treat *ρ*_*G*_ as a parameter, effectively decoupling the system.

The partition functions, *Q*_*G*_ and *Q*_*R*_, contains information about the possible states of the system when the virus is bound to either the surface or the HS chains respectively. We can calculate the partition function taking the product of the individual contributions to the respective partition function.

For *Q*_*G*_, we have contributions from the HS binding, 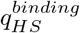 as well as a contribution from the steric repulsion arising from insertion into the chain, 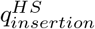,

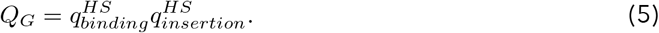

Likewise, for *Q*_*R*_ we have contributions from the receptor binding 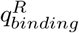, as well as the steric repulsion generated by the surrounding HS 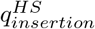,

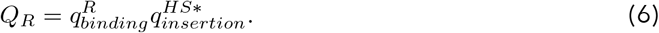

It should be noted that 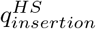 and 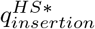 are slightly different, due to differences in the geometry of binding in the different cases. The individual partition functions will be discussed in more detail in the following subsections.

### S1 HS Binding

Each HS chain contains multiple sites to which the spike can bind. The number of these binding sites, *N*_*L*_ is some fraction of the number of units, *N*_*p*_. The number of units can be calculated from the HS chain length, *d*_*HS*_ and HS Kuhn length, *b*_*HS*_ as *N*_*p*_ = (*d*_*HS*_*b*_*HS*_)^2^.

Only a single bond can form between a given HS chain and the virus. This is likely due to the mechanical properties of the HS chain, suggesting the entropic cost of forming an additional bond is too high. We can, therefore, state the partition function for a single HS chain binding to the virus in terms of the number of spikes available for binding, *N*_*S*_, and the HS-spike binding energy in solution *ϵ*_*HS*_ :

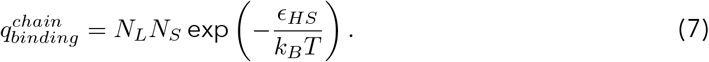

Then, we can calculate the total partition function for HS-spike binding from the binding energy of a given chain, 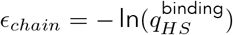, and the number of chains, as [**?**]

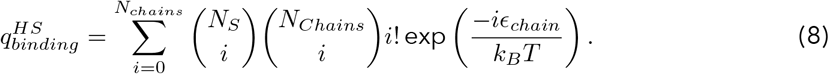

#### S1 HS Repulsion

As multiple HS chains will bind to a single proteoglycan, we expect the density of the HS chains to be non-uniformly distributed, with areas of locally high density around each proteoglycan. This means when the virus is bound only to HS chains, there will be effectively a high local density then when the virus is bound to the receptors.

We, therefore, have to consider two cases for the steric repulsion. First, when the virus is bound only to HS chains, and the second when the virus is bound to the receptors. For the receptor binding case, we will take the global HS density as *ρ*_*HS*_, while for the HS binding case we will take the higher local density.

We can then calculate the energy of insertion in terms of the density of chains, the inserted volume, *V*_*µ*_(*z*), and the insertion parameter,*δ* This gives

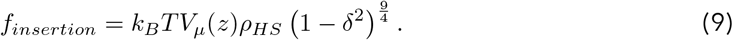

From this expression, we can calculate the partition function as 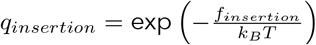.

The inserted volume can be calculated geometrically by considering the distance from the surface, *z*, as

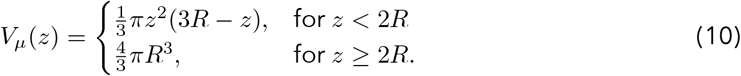

#### Insertion Parameter

The insertion parameter quantifies how far into the brush the virus has penetrated, and is given as the ratio between the brush length and the distance between the virus and surface. It can be expressed in terms of the HS chain length, *d*_*HS*_, the distance between the virus and the cell surface, *z*, and the receptor tether length, *d*_*r*_, as

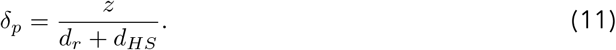

In the presence of receptor-spike binding, the cell-virus distance will be equal to the receptor tether length, *d*_*r*_. However, in the absence of receptor-spike binding, situation is more complex. We will number the binding sites of a HS chain, from *n* = 1, …, *N*_*L*_, where *n* = 1 is the site farthest from the cell surface. We can then define the distance between two sites, *n* and *n*+1, as *d*_*n*_. It is not possible to identify each distance between sites, so we will assume that the distances can be approximated by the average distance, an assumption that holds for long chain lengths with roughly equal distribution of sites. We can then express the inter-site distance as

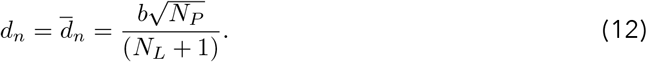

The virus-cell distance can then be expressed as a function of the closest binding site to the surface, *n*_*max*_;

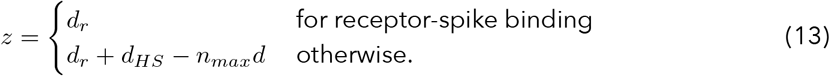

#### Mean Field Approximation

As we cannot know the distance of each virus from the cell surface, we will apply a mean field approximation, treating the repulsion as the repulsion at the average distance.

To do this, we must find the probability that a given site, *n*, is the closest bound site to the surface on a given chain. This can be expressed in terms of the probability that no bonds are formed on any HS chain at a given site, 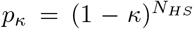, where *κ* is the probability a given site is bound (which can be calculated from the binding energy of a given site as described in equation (19))

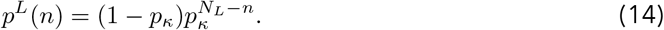

The expectation value of the cell-virus distance is then given by

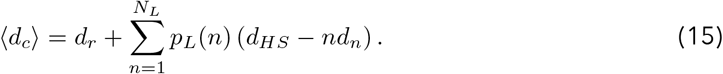

### S1 Receptor Interaction

As there are multiple possible receptors that the spike can bind to, we need to take a different but equivalent approach for calculating the receptor-spike partition function. First, we define the partition function as

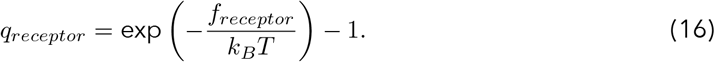

Then, we can define the energy of binding in terms of the probability that a given site is unbound, *p*_*i*_.

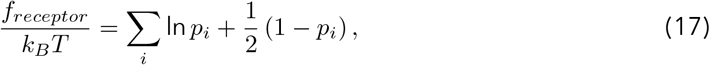

The probability of a linker being unbound can be cacluated by closure (i.e. all probabilities must sum to 1). Defining *p*_*ij*_ as the probability linker *i* is bound to linker *j*, we have

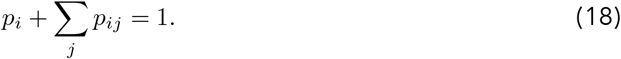

Finally, we can calculate *p*_*ij*_ can be defined as the product of the probability that both *i* and *j* are unbound and the probability that a bond will form, which is given by the Boltzmann factor – the negative exponential of the binding energy. This gives the expression

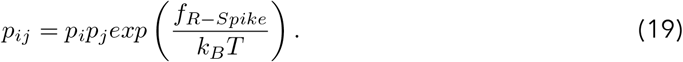

Where *f*_*R-Spike*_ is the Receptor-Spike binding energy.

#### Receptor-Spike Binding Energy

The binding energy between the spike and a given receptor has both an attractive component, arising from the binding interaction, and a repulsive component, arising from the steric repulsion caused by trapped glycans, as discussed in the main body.

As the spike may have multiple binding sites for a given receptor, we can calculate the receptor-spike binding energy in terms of the number of binding sites, *N*_*Site*_, the binding energy for a single site, *ϵ*_*R*_, and the repulsive contribution *u*_*R*_, as

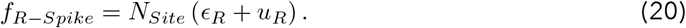

For TMPRSS2 we have only a single binding site, and we do not see the presence of glycans upon binding. Therefore, the TMPRSS2-Spike binding energy simplifies to 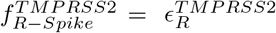.ACE2, meanwhile, has three binding sites, and we expect to see the presence of glycans upon binding. Therefore we need to calculate the repulsive contribution form.

We start by considering the Kuhn length of the glycans, *b*_*G*_; the unit size at which each unit can freely move, and orient in any direction. All details of chemical arrangements, bond rotation constraints, monomer-monomer interaction and monomer interaction with its local environment, are described by this parameter. A larger Kuhn length corresponds to a more rigid, and therefore less coiled, chain.

The mean squared end-to-end distance of a coiled polymer is then given by

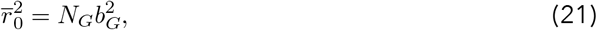

while the length of a fully extended chain is given by *r*_*max*_ = *N*_*G*_*b*_*G*_.

The probability that the end-to-end distance of a glycan is a given value, *r*, can be determined from a Gaussian distribution,

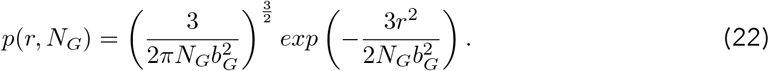

Each branch can explore a hemispherical volume of radius *r*. Therefore, integrating equation (22) over this volume gives the partition function for a given branch,

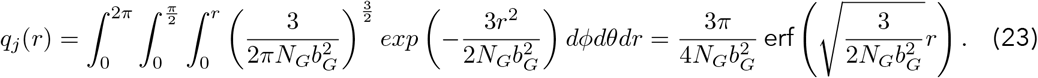

Hence the energy loss associated with the compression of a glycan with *B* branches upon binding is determined by the change in *r* from the average chain length 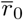 to the binding distance 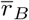,

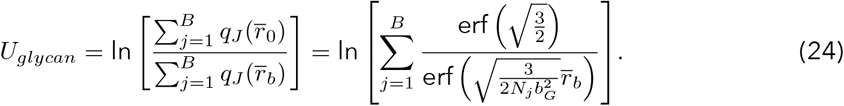

The total free energy of repulsion can now be found by summing over each glycan, 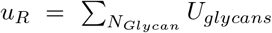, where *N*_*Glycan*_ is the number of glycans. Assuming that each glycan contains a similar amount of units, we can approximate

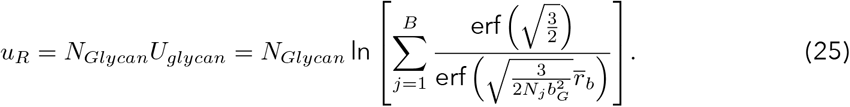

## Notes

### Competing Interest Statement

The authors have declared no competing interest.

